# Latitudinal clines in the phenology of floral display associated with adaptive evolution during a biological invasion

**DOI:** 10.1101/2024.05.30.596697

**Authors:** Mia N. Akbar, Dale R. Moskoff, Spencer C.H. Barrett, Robert I. Colautti

## Abstract

**Premise:** Flowering phenology strongly influences reproductive success in plants. Days to first flower is easy to quantify and widely used to characterize phenology, but reproductive fitness depends on the full schedule of flower production over time.

**Methods:** We examined floral display traits associated with rapid adaptive evolution and range expansion among thirteen populations of *Lythrum salicaria*, sampled along a 10-degree latitudinal gradient in eastern North America. We grew these collections in a common garden field experiment at a mid-latitude site and quantified variation in flowering schedule shape using Principal Coordinates Analysis (PCoA) and quantitative metrics analogous to central moments of probability distributions (i.e., mean, variance, skew, and kurtosis).

**Key Results:** Consistent with earlier evidence for adaptation to shorter growing seasons, we found that populations from higher latitudes had earlier start and mean flowering day, on average, when compared to populations from southern latitudes. Flowering skew increased with latitude whereas kurtosis decreased, consistent with a bet-hedging strategy in biotic environments with more herbivores and greater competition for pollinators.

**Conclusions:** Heritable clines in flowering schedules are consistent with adaptive evolution in response to a predicted shift toward weaker biotic interactions and less variable but more stressful abiotic environments at higher latitudes, potentially contributing to rapid evolution and range expansion of this invasive species.

## INTRODUCTION

Phenology affects how organisms interact with their environment, with important consequences for their survival and reproduction. As targets of natural selection and determinants of population vital rates, the timing of growth, reproduction, and other phenological traits are crucial components of eco-evolutionary dynamics. Phenology governs interaction with mutualists (Fenster et al., 2004), antagonists (Strauss & Whittall, 2006) and abiotic factors (Jin *et al*., 2019) that can vary over space and time. Consequently, plants experience spatial and temporal variation in natural selection on flowering phenology (Kudo, 2006), sometimes resulting in the evolution of local adaptation (Anderson et al., 2012; Colautti & Barrett, 2013; Preite et al., 2015). Flowering phenology is therefore a useful composite trait for investigating evolutionary responses to past and future environmental challenges (Fitchett et al., 2015; D. W. Inouye, 2022; Panchen & Gorelick, 2017; Prather et al., 2023).

Days to first flower (i.e., flowering time) is a simple yet informative metric with which to study the ecology and evolution of phenology in plant populations. For example, latitudinal and altitudinal clines in days to first flower are commonly reported in a variety of plant taxa (Alexander et al., 2009; Colautti & Lau, 2015; Ensing & Eckert, 2019; Halbritter et al., 2018; Olsson & Ågren, 2002; Panchen, 2022), and reciprocal transplant experiments have confirmed that these clines are an important component of local adaptation (Anderson et al., 2012; Colautti & Barrett, 2013; Griffith & Watson, 2005). Flowering time clines are consistent with local adaptation models characterized by stabilizing selection within populations but a shift in optimal flowering time along environmental gradients (de Villemereuil et al., 2020; Gauzere et al., 2020; Kirkpatrick & Barton, 1997).

In contrast to the prediction of stabilizing selection, empirical measurements of selection differentials suggest that flowering time is typically under directional selection within populations (Munguía-Rosas et al., 2011). Despite evidence for consistent directional selection, long-term phenology records show a variety of responses to climate change -- from earlier to later to no change in flowering time, depending on the species (Rafferty et al., 2020; Wolkovich et al., 2012). Thus, models and empirical observations of geographic clines and local adaptation appear to disagree with observed measurements of natural selection and variable phenotypic shifts in natural populations (Austen et al., 2017). Antagonistic selection on unmeasured traits correlated with the onset of flowering could help to resolve this apparent contradiction.

Day of first flowering is but one aspect of the “flowering schedule” of individual plants -- a time-series of flower production that characterizes reproductive opportunity (CaraDonna et al., 2014; Fox, 2003; B. D. Inouye et al., 2019; Newstrom et al., 1994). Although rarely examined in studies of flowering phenology evolution, other characteristics of the flowering schedule can have important effects on reproductive fitness (Ehrlén & Valdés, 2024; Forrest & Thomson, 2010; Fox, 2003). For example, the shape of the flowering schedule can be compared using metrics analogous to central moments of probability distributions including the mean, variance, skew, and kurtosis (Box 1) and ordination methods like Principal Coordinates Analysis (PCoA; Austen et al., 2014).

### Box 1: Summary characteristics of flowering schedules

Central moments (i.e., mean, variance, skew, and kurtosis) arise from probability theory and characterize distributions that can be visualized as histograms with observed values on the x-axis and probability density, frequency, or number of observations on the y-axis. Here, we use the same equations to describe analogous characteristics of flowering schedules that arise from developmental rather than probabilistic processes. In contrast to probability histograms, flowering schedules can be visualized as a time series with number or proportion of flowers on the y-axis. Although the underlying processes are distinct, the equations are the same (Table 1). To understand the biological significance of mean, variance, skew and kurtosis, we contrast flowering schedules of different shape (Fig. 1). Standardizing to proportion of open flowers (P_t_) over time (t), rather than total flower number (N_t_) accounts for variation in total flower number per plant. Moreover, we can use proportions to calculate a weighted mean day of flowering. By analogy to the mean of a probability distribution, the weighted mean of a flowering schedule represents the ‘balance point’, which better captures the start of anthesis of a typical flower.

As described in Table 1 and visualized in Figure 1, central moments capture biologically meaningful variation in flowering schedules that may not be correlated with onset or duration of flowering. For example, the variance parameter (a^-2^) describes how much flowering is spread out, even if the start and end dates are the same. An individual’s coefficient of skew (*CS_i_*) accounts for expected differences in symmetry and thus describes whether flowering is concentrated earlier (i.e., positive skew, *CS_i_* > 0 as shown by the blue dashed line) or later in the schedule (i.e., negative skew, *CS_i_* < 0 as shown by the purple dotted line), relative to the mean flowering date. Finally, the coefficient of kurtosis (*CK_i_*) describes deviations from the Gaussian expectation, contrasting individuals with a concentrated cluster of open flowers and a few flowers over a longer timeframe (i.e., leptokurtic distribution, *CK_i_* > 0 as shown in the gold dashed line) to individuals that spread out flowering more evenly over the entire schedule (i.e., platykurtic, *CK_i_* < 0 as shown by the dashed green line).

Not to be confused with the central moments of a statistical population or sample, the analogous characteristics of a flowering schedule are not probabilistic, but rather quantify temporal variation in relative reproductive investment through time (Clark & Thompson, 2011). These schedule characteristics use mathematical equations analogous to probability moments (Table 1), but are not shaped by probabilistic process and therefore may not necessarily correlate with onset or duration of flowering (Fig. 1). However, central moments are based on mean deviations with higher-order exponents, making them sensitive to outliers and inappropriate for comparing multi-modal distributions with two or more flowering peaks. As a complementary approach, PCoA compares entire schedules, regardless of shape, but must be scaled to account for differences in onset and duration of flowering. These metrics are rarely used in studies of flowering phenology, yet they have the potential to offer a novel perspective on selection and trade-offs affecting the evolution of phenology.

**Figure 1.**
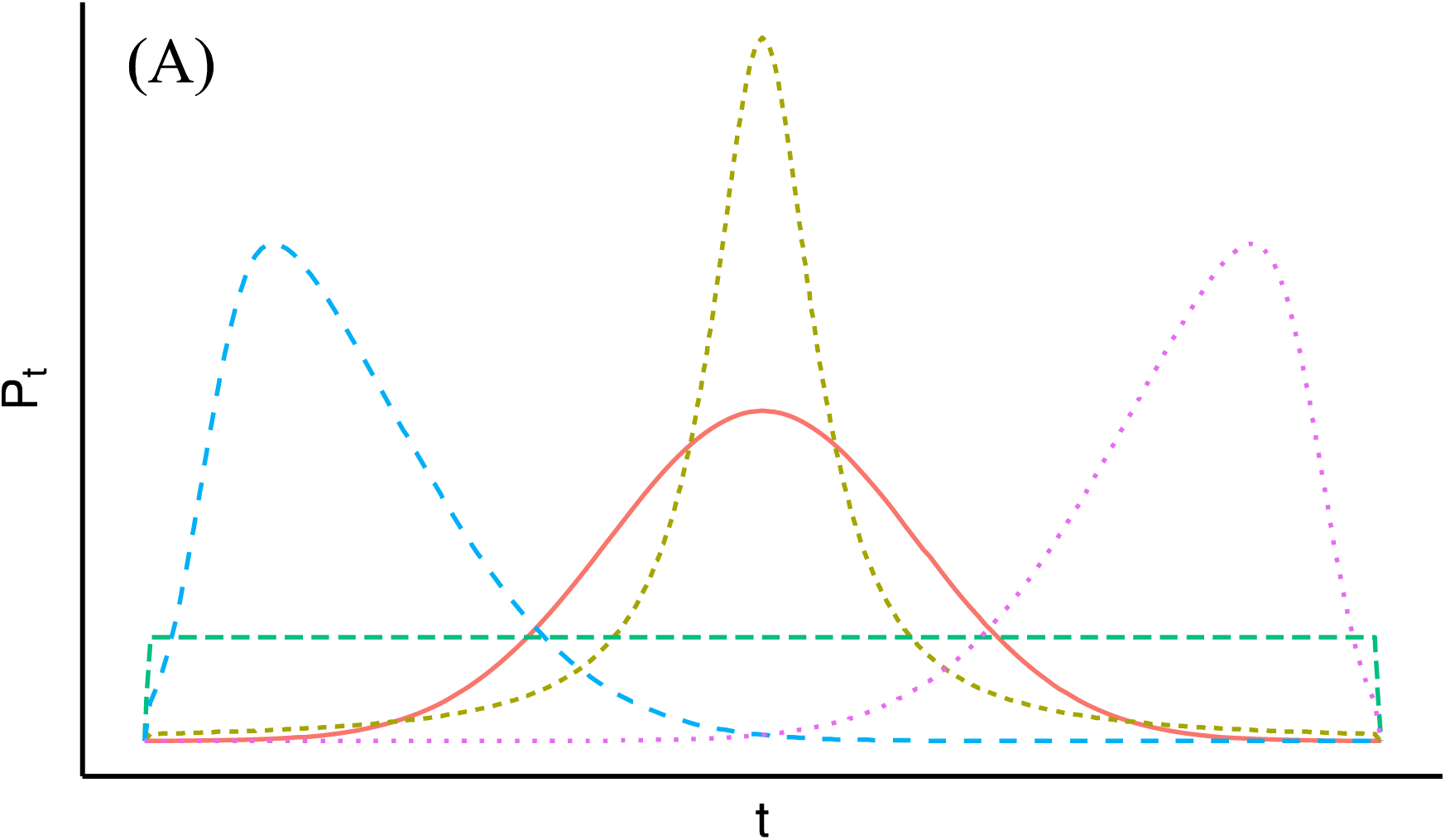
Hypothetical flowering schedules of five individuals with different characteristics, as described in Box 1. Flowering schedules are scaled to the proportion of total flower production per plant (P_t_) over time (t).

**Table 1.** Equations of average flowering schedule characteristics for a population, with subscripts *i* for individual and *t* for census time point.

| Summary Statistic | Calculation |
| --- | --- |
| Proportion ($P$) of flowers ($F$) produced at time $t$ | $P_{i,t} = \frac{F_{i,t}}{\sum_t F_{i,t}}$ |
| Day of first flower | $\alpha_i = \min\{t \mid F_{i,t} > 0\}$ |
| Day of last flower | $\Omega_i = \max\{t \mid F_{i,t} > 0\}$ |
| Duration of flowering | $\Omega_i - \alpha_i$ |
| Day of mean flowering | $\mu_i = \sum_t t P_{i,t}$ |
| Flowering schedule variance | $\sigma_i^2 = \sum_t (t - \mu)^2 P_{i,t}$ |
| Flowering schedule coefficient of skew | $CS_i = \frac{\sum_t (t - \mu_i)^3 P_{i,t}}{\sigma_i^3}$ |
| Flowering schedule coefficient of kurtosis | $CK_i = \frac{\sum_t (t - \mu_i)^4 P_{i,t}}{\sigma_i^4} - 3$ |

Central moments of individual flowering schedules describe the timing and intensity of reproductive opportunity, and thus are likely to experience strong selection from abiotic or biotic features of the environment. We are not aware of studies quantifying evolutionary divergence of entire flowering schedules among naturally occurring populations, but there are reasons to expect populations to evolve differences, as summarized in Table 2. For example, individuals that experience strong competition for pollinators may benefit from showy inflorescences that enhance pollinator attraction (O’Neil, 1997; Thomson, 1980), resulting in a more concentrated display characterized by a flowering schedule with a positive skew, smaller variance, and shorter duration. In addition, herbivores and seed predators often select for delayed reproduction (Biere & Antonovics, 1996; Elzinga et al., 2007; Juenger & Bergelson, 1998; Wright & Meagher, 2003), resulting in a later mean flowering and a negative skew in the flowering schedule. More generally, biotic interactions that are patchy and variable among years may favour a bet-hedging strategy (Elzinga et al., 2007) in which flower production is spread out over an extended period, resulting in a longer flowering duration, high flowering schedule variance, and positive kurtosis. These examples illustrate how populations experiencing different abiotic environments may evolve differences in flowering schedule traits (Table 2).

**Table 2.** Five hypotheses summarizing the effect of biotic or abiotic selection on characteristics of flowering schedules and their predicted changes with latitude (+/- refers to the predicted sign of the correlation).

| Selection Mechanism | Schedule Effect | Predicted Cline |  |  |  |  |  |
| --- | --- | --- | --- | --- | --- | --- | --- |
|  |  | Start | Duration | Mean | Variance | Skew | Kurtosis |
| *Biotic interactions* |  |  |  |  |  |  |  |
| Competition for pollinators at lower latitude | Positive skew<sup>10</sup>, shorter duration<sup>1</sup>, lower variance<sup>15,12</sup> |  | + |  | + | - |  |
| Pollinator or mate limitation at higher latitudes | Positive skew<sup>10</sup>, longer duration<sup>2,10,15</sup>, lower variance<sup>1,12</sup> |  | + |  | - | + |  |
| Herbivory and seed predation at lower latitude | Delayed start<sup>3,5,8,11</sup> and mean<sup>1,3,5,8,11</sup>, negative skew, longer duration<sup>14, 15</sup> | - | - | - |  | + |  |
| Bet-hedging to patchy biotic interactions at lower latitudes | Longer duration<sup>2</sup>, higher variance<sup>13</sup>, positive kurtosis<sup>13</sup> |  | - |  | - |  | - |
| *Abiotic factors* |  |  |  |  |  |  |  |
| Shorter season length at higher latitude | Early start<sup>4</sup>, early mean, shorter duration, smaller variance, less positive skew<sup>6</sup> | - | - | - | - | - |  |
| Bet-hedging to less predictable seasonality at higher latitudes | Longer duration<sup>2</sup>, positive kurtosis<sup>13</sup>, higher variance<sup>13</sup> |  | + |  | + |  | + |

References (Alphabetical): 1. Augspurger, 1981; 2. Austen et al., 2017; 3. Biere & Antonovics, 1996; 4. Colautti & Barrett, 2010; 5. Elzinga et al., 2007; 6. Forrest & Thomson, 2010; 7. Inouye, 2008; 8. Juenger & Bergelson, 1998; 9. Marquis, 1988; 10. O'Neil, 1997; 11. Pilon, 2000; 12. Rathcke & Lacey, 1985; 13. Simons, 2011; 14. van Doorn, 1997; 15. Wright & Meagher, 2003.

In addition to biotic interactions, flowering schedules are likely to be shaped by abiotic factors that determine the timing and duration of environmental conditions favorable to growth and development (Table 2). Growing seasons that are shortened by reduced precipitation or growing degree days may select for a more rapid flowering strategy characterized by an earlier onset of flowering, with a shorter duration and smaller variance of the flowering schedule (Austen & Weis, 2015; Colautti & Barrett, 2013; Franks & Weis, 2008; Griffith & Watson, 2005). In environments with less predictable growing conditions, it may be advantageous for plants to produce a few flowers over a longer timeframe as a bet-hedging strategy (Forrest & Thomson, 2010; Simons, 2011; Tufto, 2015), resulting in a long-tailed leptokurtic flowering schedule with higher variance.

Environmental gradients associated with latitude often covary with biotic and abiotic agents of natural selection and can be useful for studying phenotypic evolution. For example, polar latitudes typically have fewer growing degree days (reviewed in De Frenne et al., 2013), with higher seasonality and inter-annual variation (Pau et al., 2011) and weaker biotic interactions (reviewed in Zvereva & Kozlov, 2021). Adaptation along environmental gradients is commonly manifested as clines in phenotypic traits that are observable when individuals are grown in a common garden environment (Langlet, 1971; Schwinning et al., 2022). For example, adaptive flowering time clines have evolved rapidly in the perennial wetland herb *Lythrum salicaria* (purple loosestrife) during a century of invasion across North America. Selection splines (Colautti & Barrett, 2010), trait correlations (Colautti & Barrett, 2011), reciprocal transplants (Colautti & Barrett, 2013), and herbarium records (Wu & Colautti, 2022) collectively support a model of selection in which shorter growing seasons toward the northern range limit favour earlier flowering. In contrast, longer growing seasons in the southern part of the introduced range relax selection on flowering time and favor plants that flower later but grow larger in stature with more flowers. However, by focusing on the first day of flowering, previous studies of *L*. *salicaria* overlooked other characteristics of the flowering schedule, which may be important for understanding adaptation and constraint during its evolution and invasion in eastern North America.

Here, we investigate clines in flowering schedules among thirteen populations of *L*. *salicaria* sampled across ten degrees of latitude in eastern North America and grown in a common garden field study at the Koffler Scientific Reserve north of Toronto, Ontario, Canada. We used PCoA and central moments characteristics of individual flowering schedules to address the following questions: (1) Are common phenological metrics, like the onset and duration of flowering, associated with overlooked features of the flowering schedule such as the mean, variance, skew and kurtosis? (2) Are there latitudinal clines in flowering schedules that are consistent with the adaptive hypotheses outlined above and summarized in Table 2 and Figure 2? In line with these predicted adaptive changes, we expect the primary PCoA axes to reflect population differentiation in flowering schedules based on latitude of origin.

**Figure 2.**
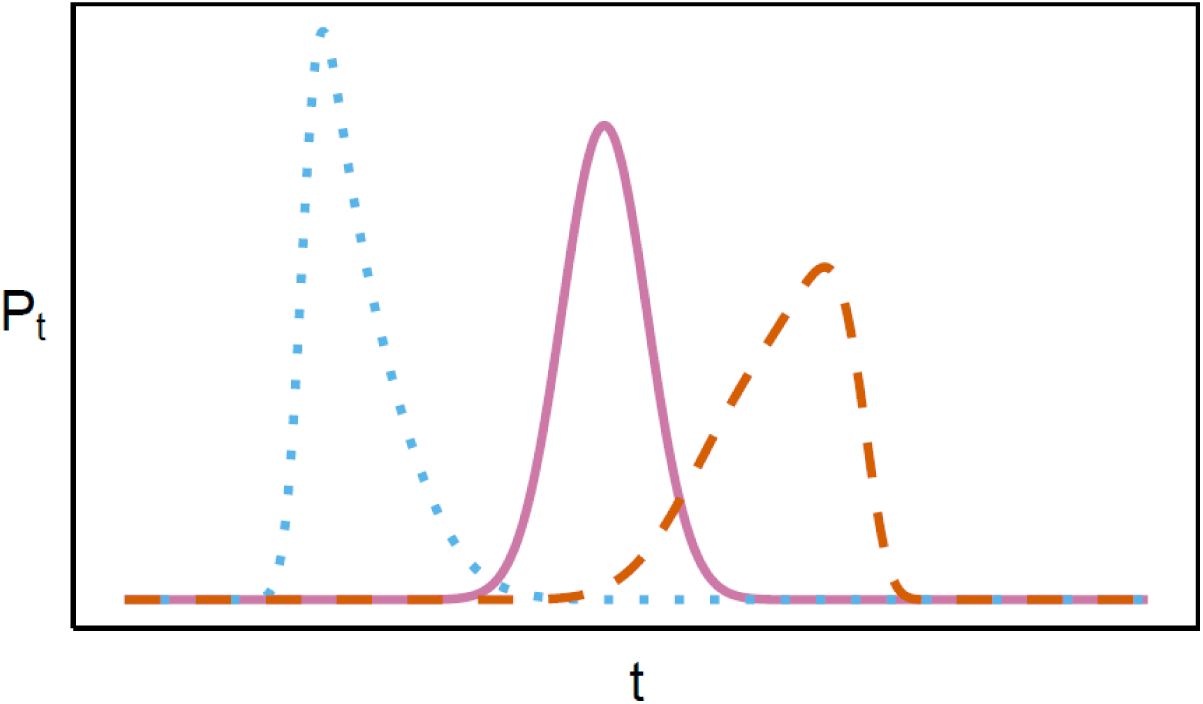
Three hypothetical flowering schedules demonstrating predicted clines in central moments from northern (blue dotted line) to southern (red dashed line) genotypes, relative to a standard normal schedule at mid-latitude (solid pink line). Each schedule is scaled as the proportion of total flowers (*P_t_*) over time (*t*). The northern schedule is defined by an earlier mean, lower variance and positive skew. The southern schedule is defined by a later mean, higher variance, negative skew, and higher kurtosis.

## METHODS

### Experimental design

Details of the field experiment are provided in Colautti & Barrett (2010) and the seeds used were collected along a 10-degree latitudinal gradient in eastern North America, as described in Montague *et al*. (2008). Briefly, populations were chosen to represent a latitudinal gradient from Timmins, Ontario (48.48 °N) to Easton, Maryland (38.75 °N) and were surveyed during the summer of 2003 (see Appendix S1 for a list of locations). When 90% of plants had finished flowering in the fall, 20-41 stems were randomly harvested from each population. Eight seeds from each of 20 seed families from each of 13 populations were grown at the Koffler Scientific Reserve at Jokers Hill, located approximately at the geographic center of the latitudinal gradient (44.03 °N). Early in the growing season (early May), plants were sprayed with insecticide to prevent the establishment of specialist beetles in the genus *Galerucella*, which were introduced for biological control of *L. salicaria*. This was sufficient to inhibit colonization of biocontrol beetles in the area, without impacting herbivore or pollinator activity throughout the growing season (May through September). The primary purpose of this experiment was to characterize flowering time clines and genetic trade-offs in a multi-year (2006-2008) common garden study, as reported in Colautti & Barrett (2010, 2011). In 2007 we additionally recorded all open flowers on every stem of a subset of 369 individuals of 216 seed families, evenly represented across the 13 populations (Appendix S1). Our daily census records of this subset of plants occurred over 125 days on a rotating schedule such that each individual was surveyed every five days from July 1 until September 30. A 5-day cycle was chosen because stigma receptivity is nearly zero after 24 h of anthesis (Waites & Ågren, 2006) and flowers are fully senescent after 5 days. Additionally, unpollinated flowers begin to senesce and abscise while pollinated flowers begin to close within 48 h of anthesis, limiting the risk of counting the same flowers on multiple census dates. All calculations and statistical analyses were conducted in R version 4.2.2 (R Core Team, 2022) in R Studio (Posit team, 2024), and are available with the original data from the Dryad repository (https://doi.org/10.5061/dryad.jdfn2z3jz).

### Comparing flowering schedules

We characterized the flowering schedules of each individual plant using the equations shown in Table 1, where the day that an open flower was first observed in the experiment is *t* = 0. Given that central moments are calculated on probabilities, we standardized the number of observed flowers for each individual on each census day as a proportion of their total number of open flowers censused over the experiment, as shown in Table 1. We then calculated a weighted census day by multiplying each day by the proportion of flowers open on that day. The proportion of flowers and the weighted census days were used to calculate the weighted mean and other characteristics of individual flowering schedules analogous to central moments (variance, skew and kurtosis) (Box 1; Table 1). Specific calculations are provided as custom R functions in the Dryad repository (https://doi.org/10.5061/dryad.jdfn2z3jz).

In addition to calculating the characteristics of every individual’s flowering schedule, we wrote a bootstrap model in R to compute a bootstrapped mean and 95% confidence intervals across 999 iterations for each population, and a similar bootstrap model to test for latitudinal clines. In each iteration, individuals were hierarchically sampled with replacement from within each population, and then the equations in Table 1 were used to calculate each schedule characteristic. The population averages of each iteration were calculated and these were used to generate 999 estimates of intercept, linear slope and quadratic regression coefficients to test for latitudinal clines. In this way we generated bootstrap means and 95% confidence intervals for each population and for the estimates of latitudinal clines. More details of the bootstrap models are available in R code from the Dryad repository (https://doi.org/10.5061/dryad.jdfn2z3jz).

In addition to the start of flowering, duration, and central moments of flowering schedules, we also used a Principal Coordinates Analysis (PCoA) to compare the entire flowering schedule shape among individuals, following methods described by Austen et al., (2014). Briefly, we computed a dissimilarity matrix from raw flower counts using Kolmogorov-Smirnov (KS) distances (Sokal & Rohlf, 1995), where each matrix cell was a KS statistic measuring the maximum distance between two flowering schedules. We chose KS distance because it is robust to differences in flower number and census schedules. However, the KS statistic is sensitive to differences in scale, so flowering durations were standardized to run from 0 to 1 in each individual flowering schedule by subtracting its earliest census day with open flowers and dividing by its total number of flowering days (see Austen et al., 2014). We used the KS matrix as the input for the **cmdscale** function to generate the PCoA axes. Since KS distances do not meet the triangle inequality condition of PCoA, any variation that is unable to be converted to Euclidean distance is represented as PCoA axes with negative eigenvalues, which may lead to overestimation of variation explained by the first axes (Austen et al., 2014; Legendre & Anderson, 1999; Podani & Miklós, 2002). Therefore, we adjusted our estimates of proportion of variation explained to only include positive eigenvalues (Austen et al., 2014). We then generated vectors to plot each flowering schedule trait in multivariate space by computing the standardized covariance of the central moments with all PCoA axes, filtered for positive eigenvalues (Legendre et al., 1998). It is important to note that the PCoA axes characterize variation in the shape of flowering schedules among individuals after controlling for variation in onset and duration of flowering. However, projecting these characteristics onto the first two PCoA axes provides a visualization of how different flowering schedule characteristics relate to the major axes of variation in schedule shape. Additionally, we colour-coded each individual by latitude of origin, to see how schedule shape varies among and within populations, and along a latitudinal gradient.

### Correlations of flowering schedule characteristics

Our first research question addresses whether conventional metrics (i.e., onset and duration of flowering) correspond to more detailed characteristics of the flowering schedule (i.e., mean, variance, skew, kurtosis, and PCoA axes). To investigate these relations, we calculated pairwise Pearson correlations to compare two scales of analysis, namely (A) average schedules of each population (*n* = 13), and (B) the deviation of each individual from its population average (*n* = 369). In addition to comparing conventional metrics with other flowering schedule characteristics, the correlational structure of these matrices can also be instructive for testing evolutionary hypotheses. For example, the evolutionary divergence of flowering schedules along biotic and abiotic gradients should produce stronger correlations among population when compared to correlations among individuals within populations.

## RESULTS

### Flowering schedule characteristics

Pearson product-moment correlations revealed significant relations among commonly measured traits of the flowering schedule (i.e., start and duration) and more detailed characteristics (i.e., mean, variance, skew, and kurtosis), both among and within populations (Fig. 3). There was no significant correlation between days to flower and duration or flowering variance among population average flowering schedules. However, there was a significant negative correlation for start date with skew and a positive correlation with kurtosis, indicating that populations with a later start date, on average, tended to have individuals with flowering clustered at later dates (Fig. 3A).

**Figure 3.**
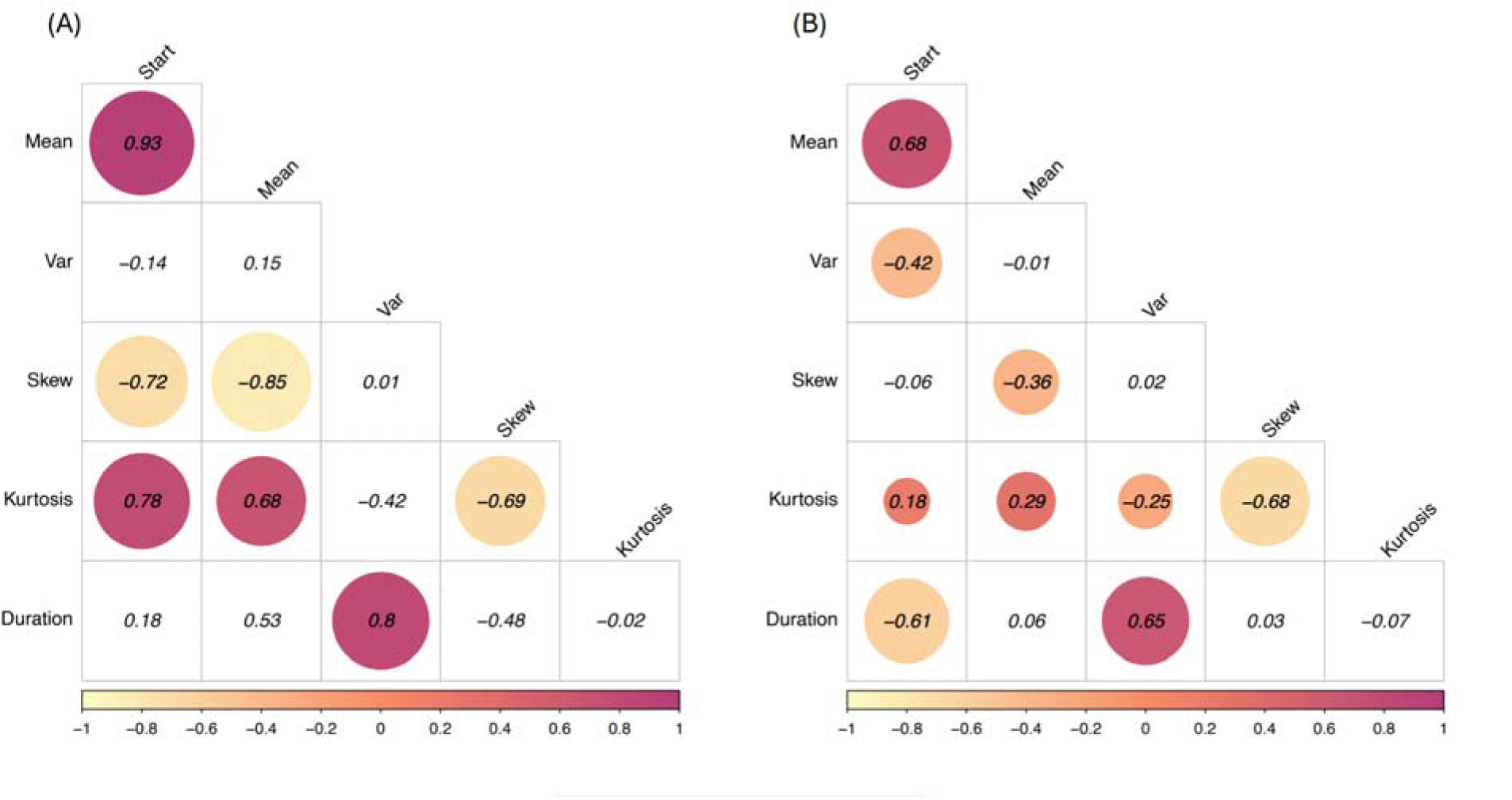
Matrices showing pairwise Pearson correlation coefficients among the start, duration, central moments (i.e., variance, skew and kurtosis), using either: (A) flowering schedule characteristics averaged across individuals of *Lythrum salicaria* within each population (*n* = 13), or (B) the residual correlations calculated on deviations of individuals from their population means (*n* = 369). Circles are shown for statistically significant correlations (*p* < 0.05), colour-coded by correlation coefficient (i.e., -1 to 1 as shown on the x-axis).

For individual flowering schedules, residual within-population correlations were generally weaker when compared to among-population correlations (Fig. 3B). However, there were negative correlations between days to flower and duration and variance of flowing among individuals within populations (Fig. 3B). These correlations were notably weaker and not statistically significant among population means (Fig. 3A). Skew and kurtosis were strongly negatively correlated both among (Fig. 3A) and within populations (Fig. 3B).

### Principal Coordinates Analysis

In addition to the above characteristics, we used Principal Coordinates Analysis (PCoA) to characterize similarities and differences among flowering schedules after controlling for variation in start date and duration. The PCoA of all 369 individuals resulted in 369 independent eigenvectors, of which 192 were positive. After adjusting the proportion of variation to include only positive eigenvalues, only the first two of 192 PCoA axes had eigenvectors that explained more than 10% of the variation in flowing schedule shape, and together they explained 35% of the variation (Fig. 4). By comparison, the next ten eigenvectors combined explained another 35% of the variation, with the remaining 180 positive eigenvectors accounting for 30% of the variation. Thus, we focus here on interpreting the first two eigenvectors. The first axis (PCoA1) was negatively correlated with mean days to flower, even though PCoA1 controls for variation in onset and duration of flowering (Fig. 4). In contrast, higher values of PCoA2 were primarily associated with higher flowering variance and lower kurtosis in flowering schedules. Despite standardizing individual flowering schedules to have the same onset and duration of flowering prior to PCoA, we found that the start and duration of flowering remained correlated with PCoA1 and PCoA2, indicating that central moments are correlated with other, unmeasured shape characteristics.

**Figure 4.**
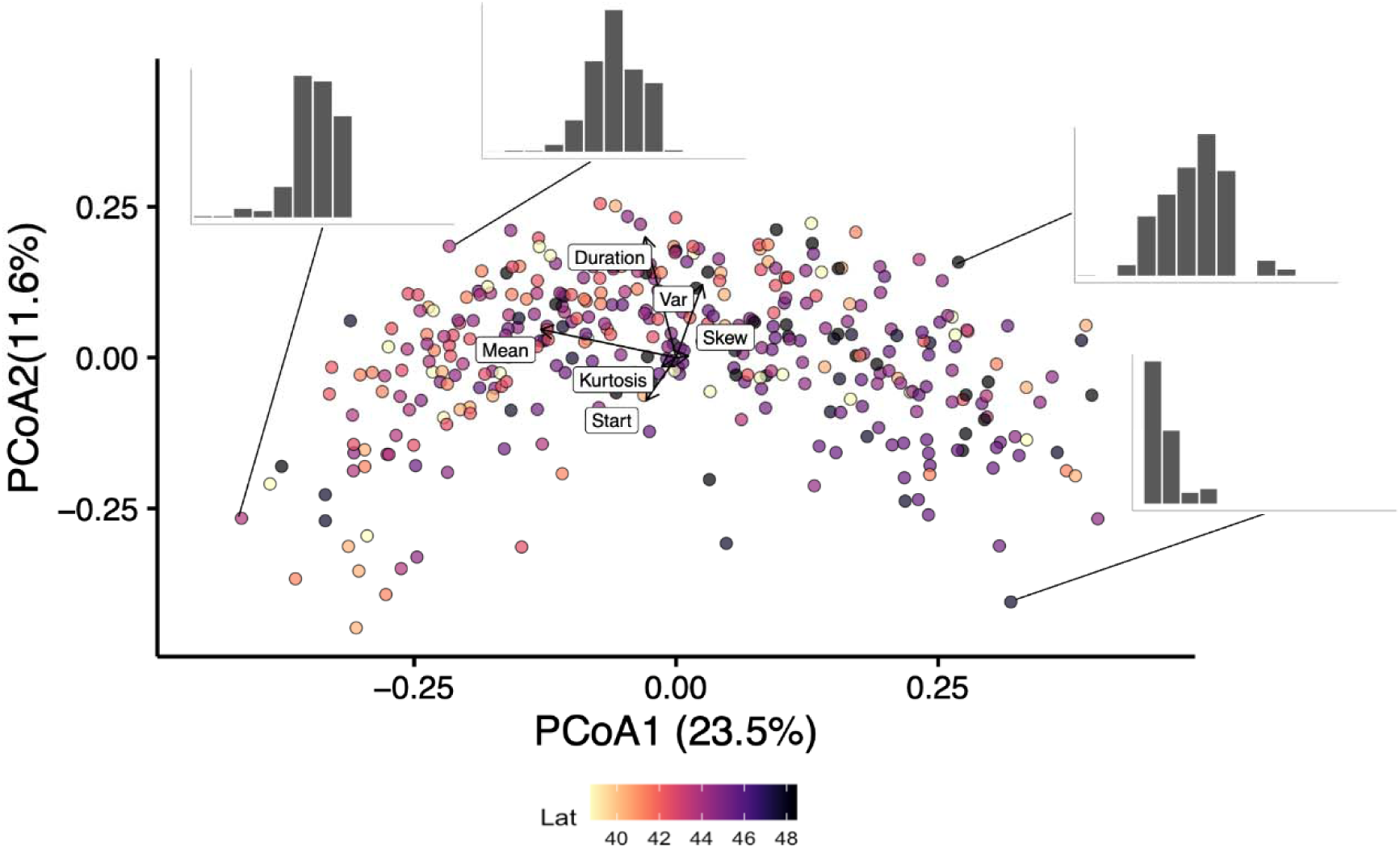
Principal Coordinates Analysis (PCoA) of Kolmogorov-Smirnov distances between flowering schedules of 369 plants from 13 populations of *Lythrum salicaria* sampled along a latitudinal gradient and grown in a common garden at the Koffler Scientific Reserve in Newmarket, ON. Point colours correspond to latitude of origin ranging from dark (48° N) to light (38° N). Bar plots show representative flowering schedules of four individual plants. Vector directions correlate to loadings on PCoA1 and PCoA2.

### Latitudinal clines

The significant correlations that we detected among populations in our central moments and PCoA analyses also covaried with latitude. For example, the start, duration, mean, and kurtosis of flowering schedules were each significantly negatively correlated with latitude (Fig. 5). This means that on average, individuals from southern populations start flowering later, have later mean flowering days, and more ‘outlier’ flowers opening far from the mean flowering date, when compared to northern populations. Additionally, average skew was negative for all populations, but more negative at lower latitudes, whereas kurtosis was close to zero but slightly positive in southern populations (Fig. 5). Thus, individuals from southern populations tended to produce more flowers far from peak flowering whereas flowering schedules in individuals from northern populations are more symmetric and concentrated (Fig. 5), even though there were no significant latitudinal clines in schedule variance.

**Figure 5.**
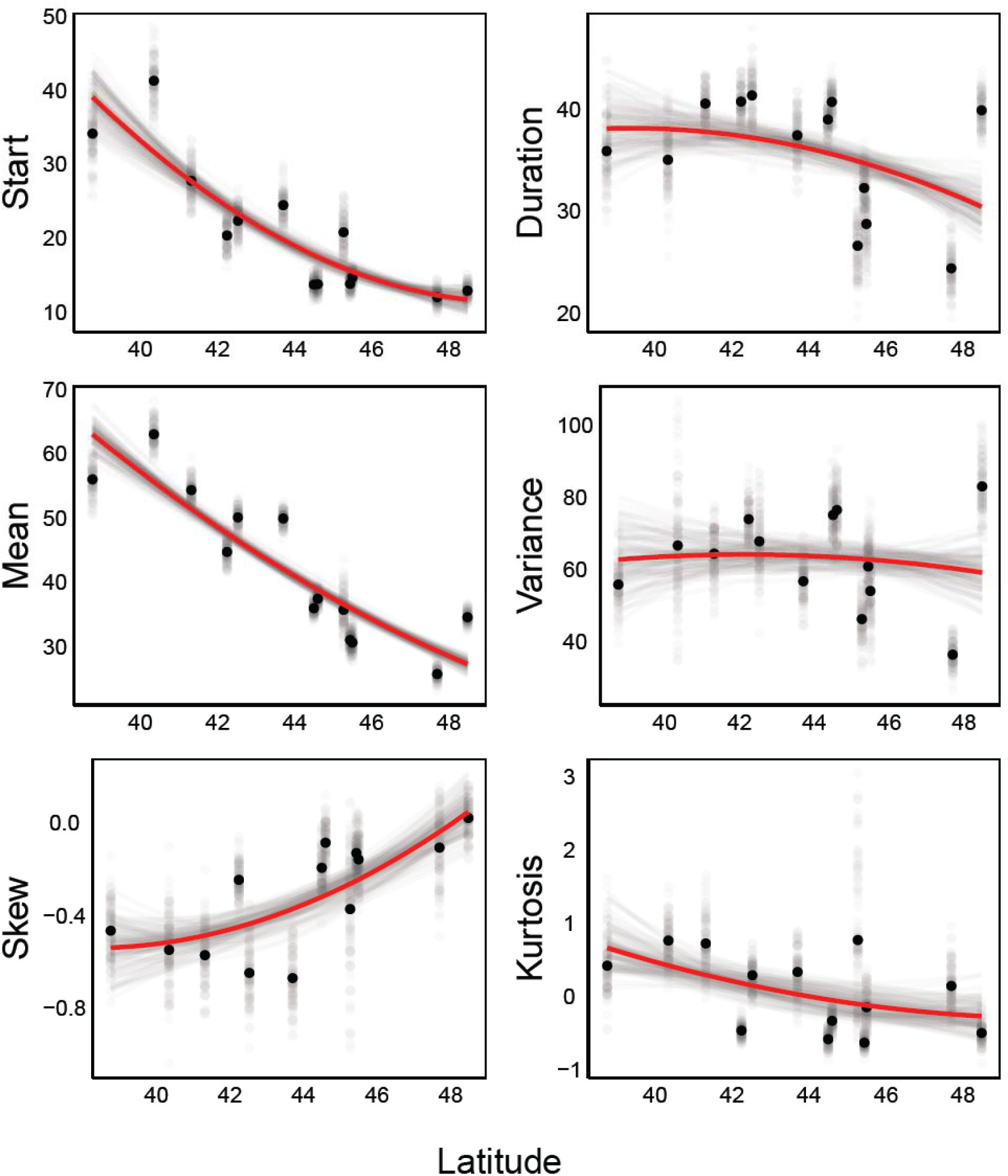
Bivariate plots showing bootstrap means and statistical relations between latitude and average flowering schedule characteristics of 13 populations of *Lythrum salicaria*, grown in a common garden field experiment at mid-latitude (44.03°N). The bootstrapped average population means (black dots) and quadratic regression lines (red curves) are based on 999 bootstrap iterations. To visualize variation among bootstrap iterations, individual iteration averages (grey dots) and regression lines (grey curves) are also shown for a random sample of 99 bootstrap iterations.

## DISCUSSION

There is growing evidence for rapid evolution in invasive species (reviewed in Barrett et al. 2017), but it remains uncertain how often adaptive evolution contributes to their spread. Previous studies focusing on the onset of flowering provided only a partial view of how plants allocate flowering resources (Box 1), leaving uncertainty about the extent to which flowering phenology is adaptive. To address this knowledge gap, we used a common garden study to test for latitudinal clines in complete flowering schedules in the wetland invader *Lythrum salicaria*. We found that the onset of flowering was significantly correlated with other aspects of the flowering schedule, both among populations (Fig. 3A), and among individuals within populations (Fig. 3B). Furthermore, the significant among-population correlations were largely consistent with evolution along latitudinal clines (Fig. 5). Compared to northern genotypes, plants from lower latitudes flowered later, for a longer duration, and with a more negative skew and leptokurtic shape (Fig. 5). Below we discuss the implications of these results for understanding the evolutionary causes and ecological consequences of flowering schedule diversity.

### Onset and duration of flowering

Date of first flower and duration of flowering are relatively easy to measure and are commonly reported as focal traits in phenological studies. However, the reproductive fitness of individuals depends on the full schedule of flower production, and it has not been clear how well the onset or duration of flowering represents other flowering schedule characteristics. In our study, the use of PCoA revealed substantial variation in flowering schedules even after standardizing for the onset and duration of flowering (Fig. 4). However, correlations of both PCoA1 and PCoA2 with onset and duration of flowering (Fig. 4) indicated that multi-dimensional variation in schedule covaries with commonly measured parameters. In line with this conclusion, the onset of flowering was strongly correlated with the mean flowering day, both among populations (Fig. 3A) and among individuals within populations (Fig. 3B). Therefore, the onset of flowering appears to be a good predictor of peak flowering in this study system.

Within populations, one might predict a trade-off between the onset and duration of flowering. Instead, we found that early flowering plants also flowered for a longer duration with a higher flowering variance (Fig. 3B). Although counterintuitive, this result is consistent with studies of other species that have reported that early flowering plants also flower for a longer duration (Austen et al., 2017; Hendry & Day, 2005). Variation in microhabitat quality (e.g., nutrient flux) and/or segregation of deleterious alleles within populations (e.g., inbreeding depression) could account for this correlation, with early/longer flowering plants representing microenvironments or genotypes of higher quality. The segregation of deleterious alleles and microhabitat variation add residual phenotypic variation that may obscure trade offs that affect clines in phenology and flowering schedules.

### Plasticity and local adaptation

Flowering schedule characteristics of individual plants can vary across years (Ehrlén & Valdés, 2024), presumably due to differences in growing environment. Previous studies of *L. salicaria* reported significant plasticity in phenology across growing environments. However, genotype-by-environment effects were relatively limited such that clines in flowering time were replicated in multiple experimental contexts, including (i) three common garden field sites spanning ten degrees of latitude (Colautti & Barrett 2013), (ii) glasshouse experiments conducted by separate laboratories (Montague et al. 2008, Colautti et al. 2010), (iii) three growing seasons in a natural field experiment (Colautti & Barrett 2010), and (iv) a ‘virtual common garden’ analysis of 3,429 herbarium specimens (Wu & Colautti 2022). These results confirm a strong heritable basis for latitudinal clines in the onset of flowering with limited genotype-by-environment interactions. However, this this may not be true for other flowering characteristics, as noted in the previous section. In particular, the mid-latitude location of our common garden site could have affected the flowering schedules of maladapted northern and southern populations.

In contrast to the highly significant relations observed between onset, duration, and variance of flowering within populations (Fig. 3B), the same correlations were non-significant among populations (Fig. 3A). Instead, duration and variance were not significantly correlated with latitude because populations originating from latitudes that were geographically closer to the common garden site tended to have longer durations and higher variance. This result is consistent with previously proposed models of constrained adaptation in this species (Colautti, Eckert, et al., 2010; Colautti & Barrett, 2011, 2013; Wu & Colautti, 2022). Specifically, populations that are locally adapted to the intermediate season length of our field site balance a trade-off between early flowering and larger growth, thus maximizing flowering duration and variance. In contrast, populations from more distant latitudes have shorter flowering durations and variance, albeit for different reasons: Early flowering plants from higher latitudes are smaller and more likely to run out of resources for flower production earlier in the season, whereas larger plants from the south have their flowering schedules truncated by the end of the growing season. This artificial truncation could also explain the observed latitudinal cline of more negative skew in plants from southern populations (Fig. 5) and the population-level correlation between days to flower and skew (Fig. 3A). Future experiments replicating common gardens at higher and lower latitudes would be necessary to test for a predicted shift from a positive cline in duration and variance at southern sites to a negative cline at higher latitudes with shorter growing seasons.

In addition to season length, the presence of specialist *Galerucella* beetles are likely to alter flowering schedules of *L*. *salicaria*, but it is unclear how this might affect the clines we observed. Released for biological control purposes, these beetles can have large effects on seed production (Grevstad 2006). More generally, herbivory can delay both the onset and peak of flowering time (Elzinga et al., 2007; Juenger & Bergelson, 1998; Pilson, 2000). A strong interaction between herbivore and host phenology might alter the clines we observed, but quantifying these effects would require a replicated common garden experiment with different levels of herbivory.

### Latitudinal clines and correlated trait evolution

If correlated clines (Fig. 3A, Fig. 5) were caused by statistical artifacts or genetic drift acting on correlated traits, then we would expect to see similar correlations among individuals within populations. Instead, we found four correlations that were much stronger among (*r*^2^ > 0.46) than within (*r*^2^ < 0.13) populations, namely start vs skew, start vs kurtosis, mean vs skew and mean vs kurtosis (Fig. 3). Without replication among individuals within families, we cannot completely rule out the possibility that environmental effects mask genetic correlations within populations (Price, Kirkpatrick and Arnold, 1988; Rausher, 1992). Nor can we completely rule out the potential role of stochastic processes without replicated sampling from independent geographical gradients (Colautti and Lau, 2015). However, the fact that flowering schedules change continuously with latitude lends additional evidence for the correlated evolution hypothesis because the direction of the observed clines are consistent with adaptive responses to abiotic and biotic selection.

Latitudinal clines in the onset of flowering have evolved independently in several unrelated angiosperm species across North America (McGoey et al., 2020; Montague et al., 2008; Samis et al., 2012; Wu & Colautti, 2022). In *L. salicaria*, we found latitudinal clines in mean, duration, skew and kurtosis of flowering schedules (Fig. 5). This latitudinal differentiation is also reflected in the PCoA (Fig. 4), wherein northern populations cluster at intermediate values of PCoA1 and higher values of PCoA2 compared to mid-latitude (higher PCoA1 but lower PCoA2) and southern populations (lower PCoA1 and lower PCoA2). These specific clines indicate that individuals from southern environments have floral displays that peak later in the season but also produce a few flowers far from the mean flowering date.

The observed clines for schedule duration, skew, and kurtosis are not consistent with bet-hedging to less predictable abiotic conditions at higher latitudes, but they do support predictions of increased intensity of biotic interactions at lower latitudes, with more variable biotic interactions at a higher intensity in southern environments (Table 2). Several effective biological control insects have been released into eastern North America, particularly *Gallerucella* beetles that feed on meristem tissue as larvae and can drastically reduce flower production (Blossey & Notzold, 1995; Grevstad, 2006; Russell-Mercier & Sargent, 2015). Previous meta-analyses have found increasing biotic interactions at lower latitudes (Zvereva & Kozlov, 2021) and towards warmer range limits (Paquette & Hargreaves, 2021), but we are not aware of prior studies showing evolved clines in flowering schedules that correspond to biotic gradients.

The negative cline in duration and the non-significant cline in variance (Fig. 5) were not consistent with the hypothesis that competition for pollinator service is more intense at lower latitudes and compatible mates are more limited at higher latitudes (Table 2). *Lythrum salicaria* is a tristylous, self-incompatible outcrosser (Colautti et al., 2010; Darwin, 1877; O’Neil, 1997) and individuals likely experience inter-specific competition for pollinators (see King & Sargent, 2012; Groulx & Sargent, 2018). A However, *L. salicaria* plants also produce larger inflorescences with more flowers at lower latitudes (Montague et al. 2008), despite increased competition for pollinators. The production of larger inflorescences with more flowers could weaken the strength of selection on flowering schedule shape because larger inflorescences are already more attractive to pollinators. Moreover, the clines predicted by gradients of mate limitation and competition for pollinators are opposite in sign to those for gradients of herbivory and seasonality. Overall, the clines we observed are most consistent with stronger selection from seasonality and herbivory than from mate limitation or interannual variation in the abiotic environment (Table 2 and Fig. 5).

### Conclusions

Our study has demonstrated that populations of *L. salicaria* in eastern North America have evolved differences in several features of their flowering schedules within a century of invasion. This phenological differentiation among populations includes predictable changes in the timing of resource allocation to inflorescence development and floral display and adds to the growing list of phenotypic traits that are targets of selection as invasive species spread along environmental gradients (MacDougall et al., 2006; McGoey et al., 2020; Molina-Montenegro et al., 2018; Urbanski et al., 2012). The ecological and evolutionary insights drawn from observations of flowering start dates alone should be interpreted cautiously, and future studies of flowering phenology should consider including features of individual flowering schedules using multivariate methods and metrics drawn from central moments theory (Table 1). These more refined characteristics of individual flowering schedules can help to evaluate adaptive hypotheses for flowering phenology and provide novel insight on the diverse ways that species respond to changing environments.

## ACKNOWLEDGEMENTS

The authors thank Z. Burivalova and R. Mackenzie for field assistance, E.J. Austen, A.E. Weis and J. Friedman for help with analysis, and E.J. Austen and an anonymous reviewer for helpful comments on the manuscript. Funding was provided from NSERC Discovery grants to S.C.H.B. and R.I.C. and NSERC CGSM to M.N.A.

## AUTHOR CONTRIBUTIONS

The conceptualization and experimental design were developed by S.C.H.B. and R.I.C, M.N.A. led the data analysis with assistance from D.R.M., S.C.H.B., and R.I.C., and all authors contributed to the manuscript, based on an original draft written by M.N.A.

## DATA AVAILABILITY STATEMENT

Raw data and fully reproducible R code for all figures, tables, and statistical analyses are available in the Dryad database (https://doi.org/10.5061/dryad.jdfn2z3jz)

---

**Appendix S1.** The distribution of 369 individuals from 216 seed families representing 13 populations of *Lythrum salicaria* sampled along a latitudinal gradient and grown in a common garden at the Koffler Scientific Reserve in Newmarket, ON, Canada.

| **Population**<br>**Latitude (°)** | **N Seed families** | **N Individuals** |
| --- | --- | --- |
| 38.75261 | 15 | 23 |
| 40.34025 | 15 | 33 |
| 41.31283 | 18 | 24 |
| 42.23822 | 20 | 34 |
| 42.52306 | 12 | 34 |
| 43.69719 | 19 | 21 |
| 44.49339 | 19 | 38 |
| 44.59489 | 19 | 22 |
| 45.26278 | 16 | 26 |
| 45.43194 | 17 | 36 |
| 45.49000 | 16 | 23 |
| 47.69081 | 14 | 32 |
| 48.47892 | 16 | 23 |

